# Utilizing Pine Needles to Temporally and Spatially Profile Per- and Polyfluoroalkyl Substances

**DOI:** 10.1101/2021.08.24.457570

**Authors:** Kaylie I. Kirkwood, Jonathon Fleming, Helen Nguyen, David M. Reif, Erin S. Baker, Scott M. Belcher

## Abstract

As concerns continue to mount over exposure to per- and polyfluoroalkyl substances (PFAS), novel methods of profiling their presence and modifications are greatly needed as some have known toxic and bioaccumulative characteristics while others have unknown effects. This task however is not simple as over 5000 PFAS of interest have been named by the Environmental Protection Agency and this list continues to grow daily. In this work, we utilized widely available archived and field-sampled pine needles and a novel non-targeted analytical method to evaluate the temporal and spatial presence of numerous PFAS. Over 70 PFAS were detected in the pine needles from this study, providing information from the last six decades related to PFAS exposure, contamination, and reduction.

---

Per- and polyfluoroalkyl substances (PFAS) are a class of manmade chemicals comprised of highly fluorinated aliphatic substances with unique anti-stick and surfactant properties. PFAS are commonly used in nonstick cookware, stain-resistant materials, food packaging, paint, and aqueous film-forming foams (AFFF). In the 1940s, the production of long-chain or legacy PFAS began in the United States (**Figure 1A**). By the early 2000s, these legacy compounds had become a global concern due to their environmental persistence, mobility, and links to adverse health outcomes including cancer, thyroid disease, immune system dysfunction, birth defects, and many others (*1*). In response, most western PFAS manufacturers have shifted to short-chain and structurally modified or emerging derivatives with ether linkages and chlorine substitutions (**Figure 1A**). While these replacements were thought to be a safe alternative to legacy PFAS, similar concerns about their environmental persistence and toxicity have also been raised (*1-6*). Thus, monitoring the presence and global distribution of this growing class of chemicals is essential.

**Figure 1.**
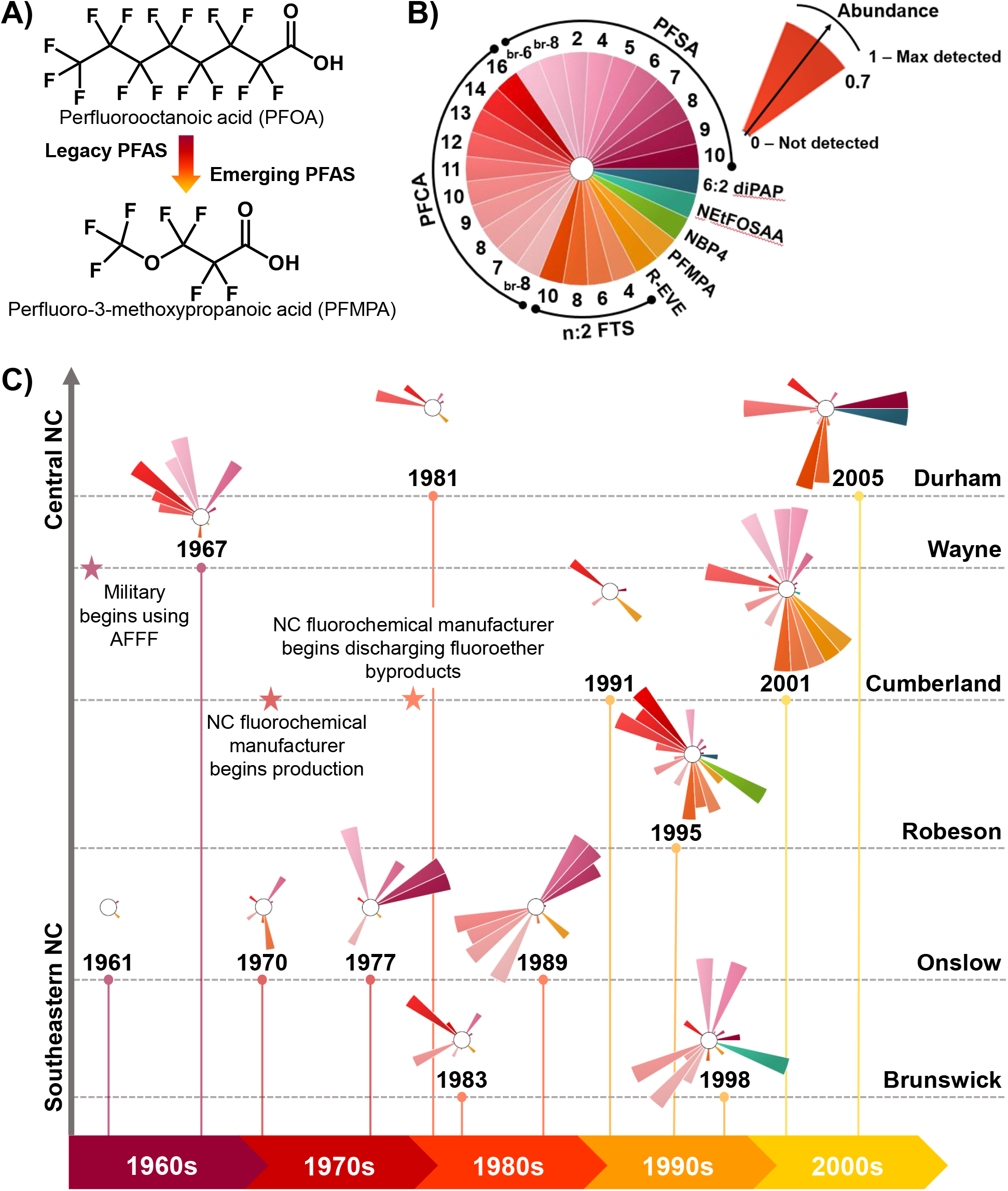
Historic North Carolina pine PFAS profiles. (**A**) Chemical structures of example legacy and emerging PFAS, perfluorooctanoic acid (PFOA) and perfluoromethoxypropionic acid (PFMPA). (**B**) Legend for the historic PFAS ToxPis in (C). Each slice represents an individual PFAS compound colored by class and the numbers indicate the numbers of carbon present in the PFAS. PFSAs, PFCAs and FTSs with two hydrocarbons (n:2) are ordered clockwise by increasing chain length with the exception of branched isomers (br-n). The height of the slice corresponds to the scaled abundance with 0 as not detected and 1 as the maximum signal detected across all samples. (**C**) Timeline visualizing the temporal and spatial PFAS trends across multiple North Caroline counties from 1961 to 2005. Each county is represented by a different row as shown on the right side of the graph. Historical information is also added on the timeline with stars to highlight some of the PFAS trends.

To date, atmospheric pollutant monitoring is particularly limited due to the expensive and complex equipment required for sampling, although it is an important route of exposure and can reflect environmental changes more quickly than lagged aquatic ecosystems (*7*). Therefore, naturally occurring passive sampling material, such as pine needles, have been utilized as an alternative for the global, low-cost sampling of a wide variety of organic pollutants, including a limited number of mostly legacy PFAS (*8-11*). Pine needles also have waxy surfaces and previous studies have shown that uptake of PFAS occurs primarily via adsorption of atmospheric PFAS to the needles with limited uptake from the soil and water. These properties enable the highly comprehensive sampling of pine trees compared to traditional monitoring techniques (*10, 11*).

Here, we explored the use of pine needles and non-targeted multidimensional mass spectrometry-based measurements for insight into legacy and emerging PFAS presence and distribution over time (**Figure S1, Tables S1-S2**) (*12*). Evaluation of all samples was performed using a non-targeted platform coupling liquid chromatography, ion mobility spectrometry, and mass spectrometry (LC-IMS-MS) separations following an optimized PFAS extraction technique which requires as few as 20 needles per sample (**Supplementary Materials, Tables S3-S6**). To extend these measurements even further, 14 heavy labeled PFAS were spiked into the extractions (**Table S7**) to add targeted quantitation capabilities to the non-targeted analyses, an approach known as simultaneous discovery and targeted monitoring (DTM), which is only possible with the multidimensional LC-IMS-MS platform (*12*). Because commonly used LC-MS methods distinguish analytes based on their hydrophobicity (retention time) and mass-to-charge ratio (*m/z*), interfacing IMS with traditional LC-MS techniques allows further separation based on the gas-phase size, shape, and charge of the ions. LC-IMS-MS measurements have also been shown to greatly enhance sensitivity in highly complex samples due to the increase peak capacity possible for the measurements (*13, 14*). While PFAS manufacturing began in the 1940s, routine testing of environmental contamination did not begin until the turn of the 21^st^ century when the toxicity and persistence of these chemicals became public knowledge. Thus, PFAS presence during this gap (1950s-1990s) has been retroactively evaluated using sediments and ice cores (*15-17*), but these methods are limited to aquatic and artic ecosystems, respectively, and may disregard atmospheric contributions and regional specificity. The LC-IMS-MS analyses of the archived pine needles in this study identified 29 unique PFAS over five decades in six North Carolina (NC) counties (**Table S8**). The resulting ToxPi profiles, which show the scaled abundance of each individual PFAS (**Figure 1B**), are displayed in **Figure 1C** for each site. While both time and location played roles in the PFAS presence, likely due to proximity to a fluorochemical manufacturer on the Cumberland-Robeson County border and other point sources such as military sites, some trends overcame regional differences. One such trend is the number of chemicals detected at each time point which is significantly different (p < 0.05) prior to and following 1985, with averages of 8 and 14 detected PFAS. This finding highlights both the increasing number of new contaminants and persistence of legacy PFAS, many of which have been phased out of use for over a decade, but are still detected globally and on a regular basis (*18*). One sample which did not fit this trend was collected in Wayne County in 1967 where 11 PFAS were detected. However, this location was near an Air Force Base in Wayne County and since aqueous film-forming foam (AFFF) containing concentrated PFAS mixtures were adopted by the military in the 1960s for fire training and emergencies, we expect this contributed to the high abundance of PFAS observed in this area (*19, 20*).

To further understand the regional trends, PFAS subclasses were also examined. The observed subclass trends appeared to be mainly due to changes in the PFAS synthesis approaches and replacement of legacy PFAS with emerging species. Starting in the 1940s, PFAS were dominantly produced via electrochemical fluorination (ECF) which yields nonspecific mixtures of even- or odd-chained and linear or branched compounds. An alternative approach that yields mostly linear species, known as fluorotelomerization, was developed in the 1970s. Fluorotelomerization has been the dominant approach since the 2000s as it is mainly used for the synthesis of short-chain emerging replacements (*21*). Overall, in **Figure 1C** the abundant perfluoroalkyl sulfonic acids (PFSAs, pink) and perfluoroalkyl carboxylic acids (PFCAs, red) shift from longer, branched chains to shorter, linear chains over time. PFAS belonging to additional subclasses then emerge following 1980, including fluorotelomer sulfonates (FTS, orange), perfluoroalkyl ether carboxylic acids (PFECAs, yellow), a perfluoroalkyl ether sulfonic acid (PFESA, green), a perfluoroalkyl sulfonamide (PFASA, aqua), and a fluorotelomer alcohol phosphate ester (blue). These findings agree with the known history of each PFAS subclass. For example, fluorotelomers, or polyfluoroalkyl substances containing aliphatic chains which are not fully fluorinated, have only been produced since the 1970s and the peak FTS levels were detected in the 2001 and 2005 needles, shortly after FTS became the principal component of AFFFs in the 2000s (*19-21*). Furthermore, the fluorochemical manufacturer in NC began production in 1971 but is not reported to have discharged fluoroether (PFECA and PFESA) byproducts into the environment until 1980 (*2*). The detected fluoroether levels increase beyond 1980 with their highest relative abundances in 1995 and 2001. Likewise, both samples with the highest fluoroether levels were sampled from the neighboring counties of the manufacturing plant. Taken together, this historical study enabled snapshots of PFAS exposure through time and illuminates an increase in PFAS complexity via rising numbers of subclasses and individual species within each subclass over time.

To gain a better understanding of PFAS spatial and temporal distribution within the last 5 years, field samples were collected from 20 locations across NC from 2017-2020 (**Figure S1, Table S1**). The IMS measurements greatly aided in the analyses of these samples as 75 PFAS were detected, 61 of which were confidently identified using a chemical standard spectral library with LC, IMS and MS information (**Tables S6, S9**). In the drift tube IMS (DTIMS) experiments, a drift tube is filled with a buffer gas which collides with ions as they traverse the tube under an applied electric field. Smaller ions have fewer collisions and migrate faster, thus they have a lower drift time. The ion’s drift time is thus directly correlated to size, known as collision cross section (CCS) (*22*). While filtration and solid-phase extraction (SPE) cleanup steps were performed prior to analysis, the biological matrix of the pine samples is quite complex, having many metabolites, lipids and other small biomolecules which leads to interferences and increased noise in traditional LC-MS analyses. These interferences impact the extracted ion abundances and subsequent quantitation and limit the number of low-abundance molecules that can be identified. In the IMS as displayed in **Figure 2A**, PFAS have much lower drift times than hydrocarbon-based biomolecules of similar mass-to-charge (*m/z*), such as lipids, due to the larger mass but similar size of fluorine compared to hydrogen. Using Skyline, an open-source software program able to match the observed multidimensional features to the LC, IMS and MS library, any signal outside of a set range of IMS drift times is filtered out of the extracted ion abundance, allowing better quantitation of the PFAS (*13*). The IMS dimension can also be leveraged to tease apart isomers that co-elute in the LC dimension, such as linear perfluorooctanesulfonic acid (PFOS) and its branched isomers such as 1-MHpS (**Figure 2B**). These constitutional isomers would both be integrated as one ion in traditional LC-MS analyses, but their near-baseline separation in the IMS drift time dimension due to the smaller size of the branched isomer allows them to be treated as two separate entities here. **Figure 2B** also demonstrates that the native ion, linear PFOS in this case, and the corresponding heavy labeled internal standard (^13^C8-PFOS) align in both the LC and IMS dimensions, allowing straightforward annotation and quantitation of PFAS with matching standards (**Table S7**). Moreover, IMS increases identification confidence due to the addition of the CCS parameter and subclass-specific trendlines when CCS is plotted as a function of *m/z* (**Figure S2**).

**Figure 2.**
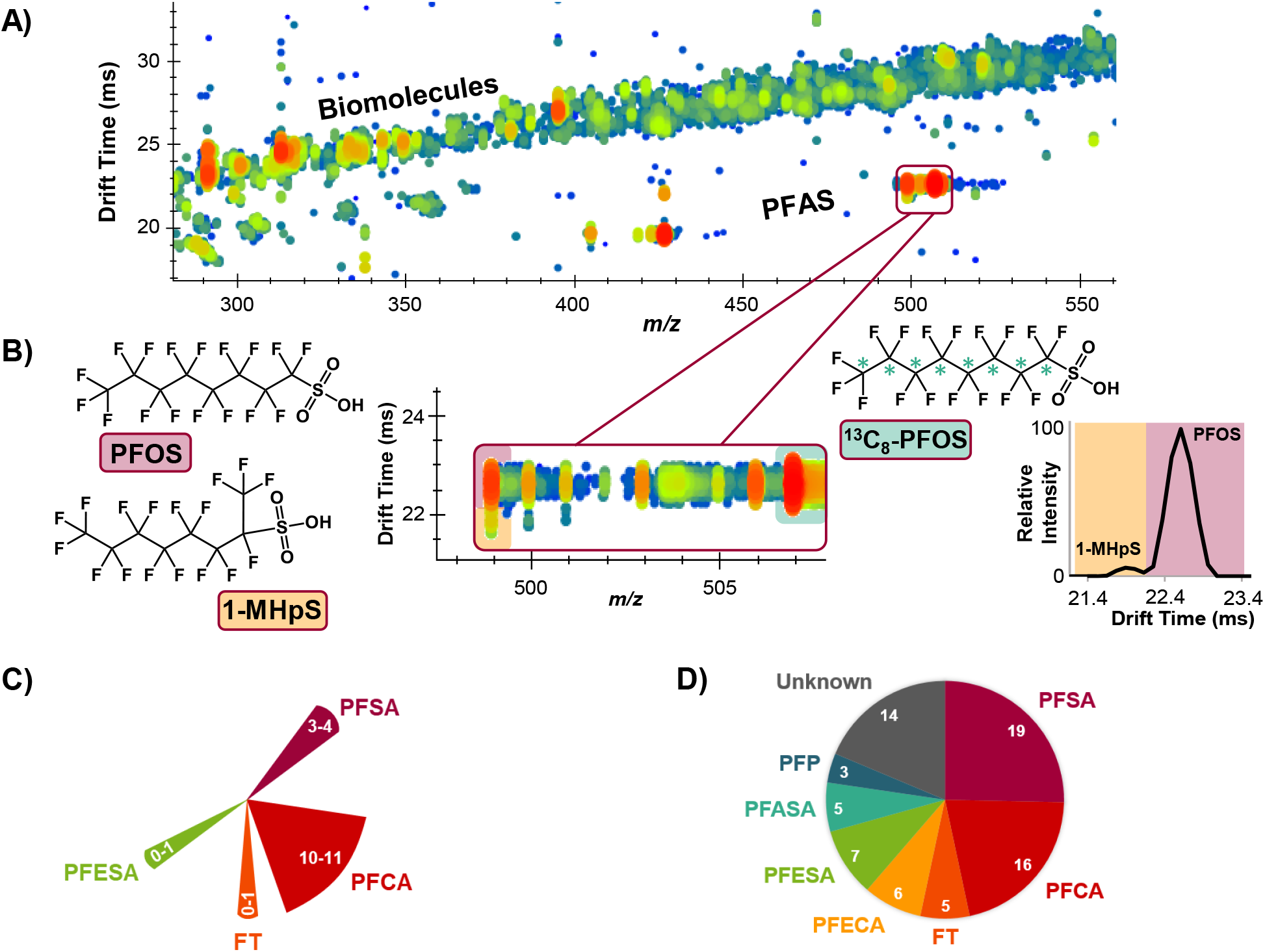
Ion mobility spectrometry capabilities. (**A**) Separation of PFAS from hydrocarbon-based biomolecules (e.g., lipids) in the IMS drift time dimension clearly distinguishes the different molecule types with similar *m/z* values. (**B**) Zoom-in of (**A**) showing the IMS drift time separation of linear PFOS (pink) and its co-eluting constitutional isomer 1-MHpS (orange), as well as the IMS drift-alignment of linear PFOS and the mass labeled linear PFOS internal standard ^13^C8-PFOS. (**C**) Breakdown of the 75 PFAS detected in the pine needles, 61 of which were confidently annotated using an LC-IMS-MS spectral library built from >100 PFAS standards.

Using this approach, a much more extensive and structurally diverse set of PFAS were identified compared to previous studies (**Figure 2C**), such as the PFSAs which range from 1-10 carbon chains and cover linear, branched, and cyclic chemicals, or the PFESAs which have varying carbon chain lengths and number of oxygen atoms as well as two chlorinated derivatives (**Table S6**). Of the 61 PFAS confidently identified in the pine needles from 2017-2020, seven structurally diverse subclasses including PFSAs, PFCAs, fluorotelomers (FTs), PFECAs, PFESAs, PFASAs, and perfluoroalkyl phosphinates/phosphate esters (PFPs) were observed (**Figure 2D**). An interactive map displaying the ToxPi profiles with all 61 identified PFAS from each field sampling location at various time points is available at https://arcg.is/19vXuK0.

To further evaluate important chemical changes occurring in the pine needle samples collected over the last five years, fluoroether distributions were investigated. These chemicals are of particular interest due to the presence of a fluorochemical manufacturer in NC which is known to discharge fluoroether byproducts into the environment, specifically the Cape Fear River, a source of drinking water for 200,000 southeastern NC residents. Moreover, fluoroethers have been detected worldwide and linked to similar health outcomes to legacy PFAS (*1, 2, 4, 6*). The amount and distribution of PFECA and PFESA chemicals in southeastern NC in 2017-2018 was compared to 2019-2020 and displayed in **Figure 3** as a ToxPi*ArcGIS map (*23*). Of the 13 detected fluoroethers, only perfluoro-3-methoxypropanoic acid (PFMPA), NVHOS, and Nafion byproduct 2 (NBP2) were detected outside of Cumberland and Robeson counties, or further than 11 miles from the manufacturing plant. The highest abundance of each compound was detected at the two sampling sites within 2 miles of the manufacturer except PFMPA, which was most abundant near the NC coast, and two chlorofluoroether sulfonic acids (Cl-PFESAs) F53-B minor (9Cl-PF3ONS) and major (11Cl-PF3OUdS), which were only detected at a nearby regional airport in 2020. The fluorochemical manufacturer has not reported production of the chlorofluoroethers, therefore these are likely a component of industrial materials at the airport. Furthermore, in December 2019, the NC fluorochemical manufacturing plant was required to install a thermal oxidizer to control PFAS emissions from the facility (*24*). This explains the drop in fluoroether abundance from 2017-2018 to 2019-2020, particularly at the sampling site approximately 2 miles from the manufacturing plant (**Figure 3**). At this site, perfluoro-2-methoxyacetic acid (PFMOAA) was no longer detected in 2019-2020, and the remaining fluoroethers, hexafluoropropylene oxide dimer acid (HFPO-DA, GenX), perfluoro-2-methoxypropanoic acid (PMPA), R-EVE, Nafion byproducts 4 and 6 (NBP4, NBP6), and NVHOS, decreased in abundance by 76-99%. While several fluoroethers at the sampling site approximately 1 mile outside of the manufacturing plant saw decreases in abundance over time, including GenX, PFMOAA, perfluoro-2-ethoxypropanoic acid (PEPA), NBP2, and NBP6, the abundance of PMPA increased and Nafion byproduct 1 (NBP1) was only detected at this location in 2019-2020. Taken together, these results show a decreasing trend in overall fluoroether abundance and spread from the manufacturing plant, except for a few species which may be recent replacements for the fluoroethers under considerable scrutiny and local regulations such as GenX (*2*). Importantly, these results demonstrate the capability of this approach for monitoring the introduction of new chemicals and tracking the success of PFAS reduction and remediation efforts, particularly due to the evident spatial and temporal resolution of this method.

**Figure 3.**
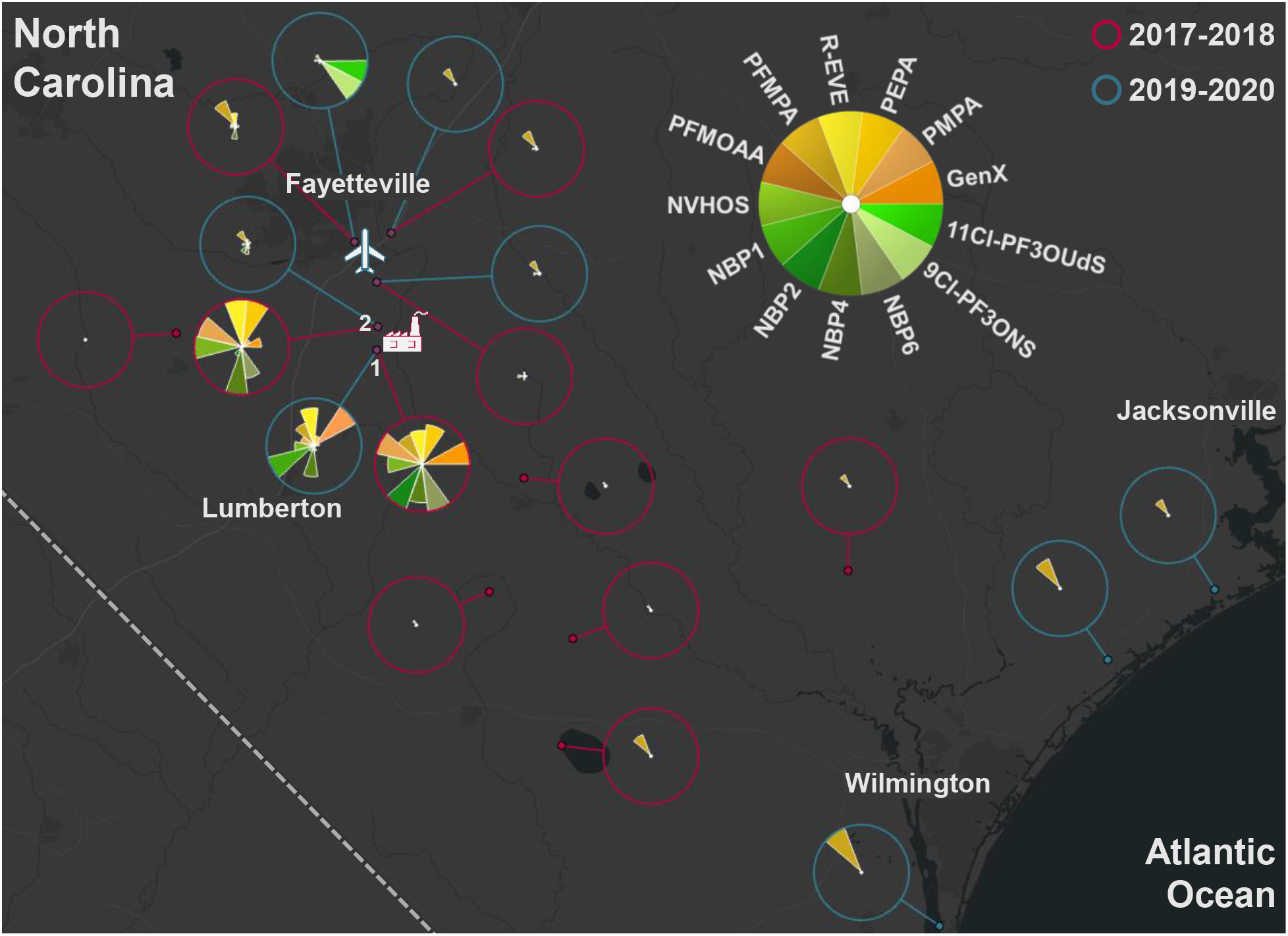
Fluoroether distribution and spread from a fluorochemical manufacturer. Map of southeastern North Carolina with averaged PFECA and PFESA ToxPi profiles of pine needles collected from 14 locations in 2017-2018 (red outer circles) and 2019-2020 (blue outer circles). The factory icon signifies the location of the fluorochemical manufacturer and the airplane icon points to the Fayetteville airport. 1 and 2 denote the sampling sites located approximately 1 and 2 miles from the fluorochemical manufacturing plant.

Beyond the fluoro24chemical manufacturing plant, other identified PFAS point sources in this study included airports and a military site. One such example is the NC National Guard within the Raleigh-Durham (RDU) International Airport area. In our study, the four sampling sites around RDU had unique ToxPi profiles, showcasing the spatial resolution and specificity of this approach, as three of the locations are barely 1 km (0.63 mi) apart. Of these four locations, the site located directly outside of the NC National Guard facility had extremely elevated levels of some PFSAs, PFCAs and PFASAs (**Figure 4**). Estimated concentrations of several PFCAs and PFSAs of interest were further examined with their corresponding internal standards (**Table S9**). Several important observations to note were that the concentrations of the linear C4 and C6 PFCAs; branched C5, C6 and C8 PFCAs; linear C4, C6 and C8 PFSAs; and branched C6 and C8 PFSAs at the National Guard sampling site ranged from approximately 100 to 90,000 times higher than the average values for the entire 2019-2020 sample set. Additionally, linear C5 PFCA; 1-MHpS; branched C4, C5, and C7 PFSAs; perfluoro-1-hexanesulfonamide (FHxSA); and two unknown PFAS were only detected at this location. These unique species, isomer profiles (**Figures S3-S4**) and elevated levels are likely due to firefighting training at the National Guard base with AFFF, which commonly contain these classes of compounds (*19, 20, 25*).

**Figure 4.**
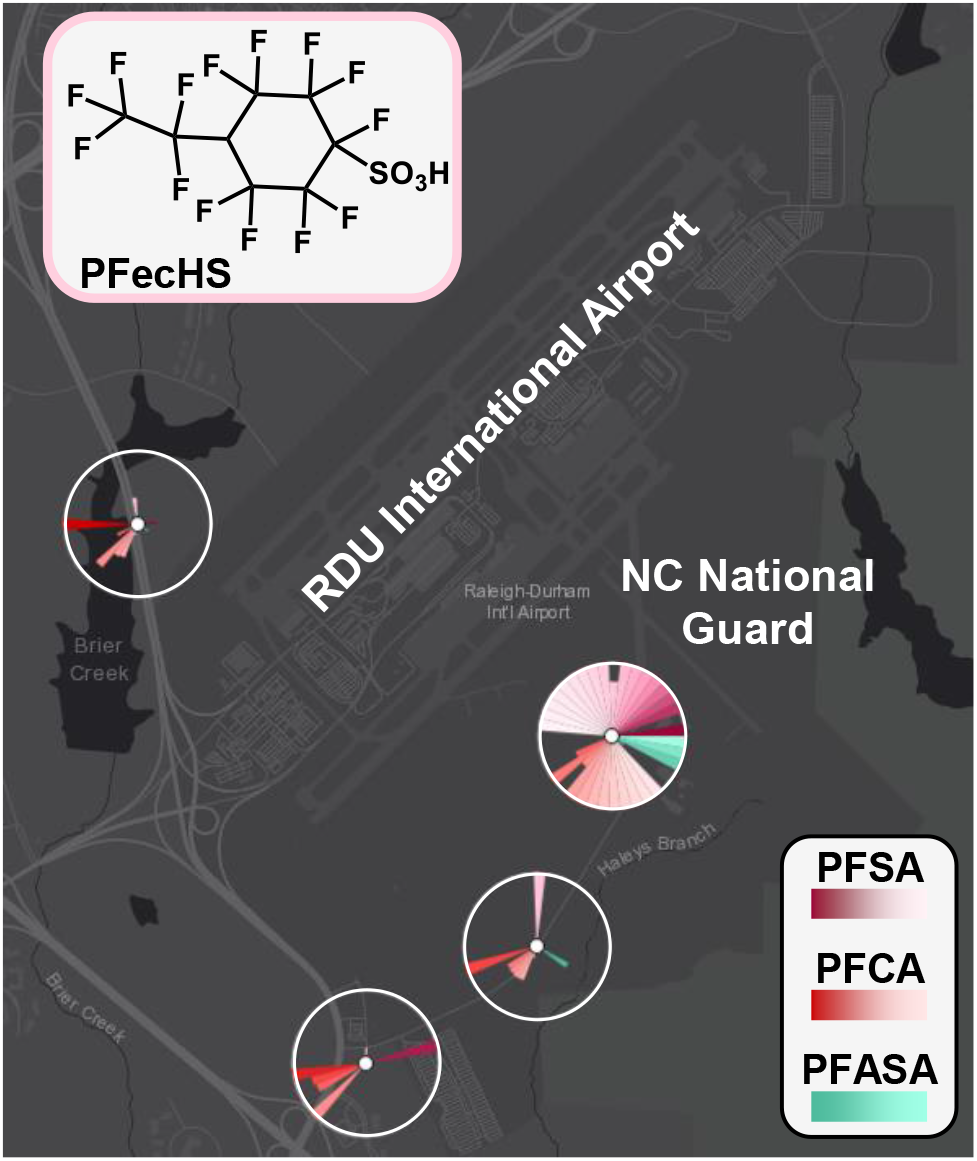
Military and international airport PFAS contributions. Map of Raleigh-Durham International Airport and the North Carolina National Guard facility with averaged ToxPi profiles of all detected PFSA, PFCA and PFASA compounds in the pine needles from four sites in 2019-2020. The chemical structure of PFECHS, a cyclic PFSA which was only detected at these four sites, is displayed.

The other three RDU International Airport sampling sites exhibited unique PFAS profiles as well, with elevated levels of long-chain PFCAs that were either not detected or detected at much lower abundance at the site located nearest the National Guard facility. Moreover, the emerging cyclic PFOS analog perfluoroethylcyclohexane sulfonate (PFECHS) was only detected at the four locations near the RDU airport (**Figure 4**). PFECHS was excluded from the previous C8-based compound restrictions due to its lack of alternatives and critical function as an erosion inhibitor in aircraft hydraulic fluids, and it continues to be detected worldwide (*26, 27*). Another emerging C8-based compound which was detected only at RDU and Fayetteville Regional (FAY) Airports in 2020 is the chlorinated derivative 8Cl-PFOS. Aside from this similarity, the FAY airport PFAS profile did not resemble those of RDU airport, as it mainly had high levels of the Cl-PFESAs as well as fluorotelomer sulfonates (**Figure S5**).

In summary, by using both archived and freshly collected pine needles in conjunction with non-targeted multidimensional analyses, we were able to monitor the spatial and temporal distribution of diverse PFAS. Our results illuminate historic PFAS profiles, identify point sources, track the introduction of new chemicals, and evaluate the success of contaminant reduction efforts. While the scope of this study was limited to North Carolina, this approach can be applied globally wherever pine trees exist to gain insight into both historic and modern PFAS profiles worldwide.

## Acknowledgements

All measurements were performed in the Molecular Education, Technology and Research Innovation Center (METRIC) at NC State University. We thank Professor Alexander Krings, Director of the NCSC Vascular Plant Herbarium and Layne Huiet, Collections Manager of Vascular Plants at the Duke University Herbarium for graciously donating archived pine needles for the historical analyses.

## Author contributions

Conceptualization: KIK, JF, DMR, ESB, SMB

Sample collection: KIK, HN, SMB

Methodology and experimental design: KIK, JF, HN, DMR, ESB, SMB

Sample preparation and LC-IMS-MS analysis: KIK, HN

Data annotation and qualitative/quantitative analysis: KIK, ESB

Data interpretation: KIK, ESB, SMB

Data visualization: KIK, JF, DMR, ESB

Development of software tools and interactive map: JF, DMR

Writing – original draft: KIK

Writing – review & editing: KIK, JF, DMR, ESB, SMB

## Data and materials availability

Skyline files with all LC-IMS-MS data are available via Panorama Public at https://panoramaweb.org/pfas_pine.url. An interactive map of the full field study dataset is also available at https://arcg.is/19vXuK0.

