## Supplementary Materials for "Utilizing Pine Needles to Temporally and Spatially Profile Per- and Polyfluoroalkyl Substances"

**This PDF file includes:**

Materials and Methods
Supplementary Text
Figs. S1 to S5
Tables S1 to S11
References

**METHODS**

Standards and Reagents

Figure S1: Pine sampling locations

Table S1: Pine field sampling locations

Table S2: Archived pine samples

Sample Collection and Treatment

LC-IMS-MS Instrumental Analysis

Table S3: LC gradient settings

Table S4: Source settings

Table S5: IMS-MS settings

Quality Assurance and Data Analysis

Table S6: PFAS LC-IMS-MS parameters

Table S7: Internal standards

**RESULTS**

Quantitative Data

Table S8: Archived pine concentration estimations

Table S9: Field sample concentration estimations

Multidimensional Data and Isomers

Figure S2: IMS-MS trendlines

Figure S3: PFCA isomeric composition

Figure S4: PFSA isomeric composition

Point Sources

Figure S5: FAY Regional Airport profile

### Materials and Methods

#### Standards and Reagents

Stable isotope labeled internal standards were obtained from Wellington Laboratories (Guelph, Canada). All experimental and quality control samples as well as extraction blanks were spiked with $^{13}C_4$-PFBA, $^{13}C_5$-PFPeA, $^{13}C_5$-PFHxA, $^{13}C_4$-PFHpA, $^{13}C_8$-PFOA, $^{13}C_9$-PFNA, $^{13}C_6$-PFDA, $^{13}C_7$-PFUdA, $^{13}C_2$-PFDoA, $^{13}C_2$-PFTeDA, $^{13}C_3$-PFBS, $^{13}C_3$-PFHxS, and $^{13}C_8$-PFOS. Additionally, archived samples and samples from location 5 (**Table S1**) were spiked with $^{13}C_3$-GenX.

For extractions and mobile phases, Optima LC-MS grade methanol, water, and ammonium acetate, ammonium hydroxide, and glacial acetic acid were obtained from Fisher Scientific.

#### Sample Collection and Treatment

Needles were collected from central and southeastern North Carolina *Pinus taeda* and *Pinus palustris* trees on public land. Sample locations are depicted in **Figure S1**. At each field sampling site, pine needles were collected from an individual tree and stored in polypropylene bags. At some sites, additional needles that had been shed the previous year were collected and stored in a separate polypropylene bag. The field samples are summarized in **Table S1**. The needle brachyblasts were separated and discarded, then the needles were incubated at 37°C for approximately 1-2 weeks to constant weight. Archived needles were obtained from either the Duke University Herbarium or the North Carolina State University (NCSC) Herbarium. These specimens were stored at room temperature within a folded paper cover since their collection date. Approximately 20 needles were transferred from each specimen to polypropylene bags following removal of the brachyblasts and stored until extraction. The archived samples are summarized in **Table S2**. The first three samples from 1938, 1947 and 1957 were discarded due to contamination from soaking the pre-1960s plant materials in various pesticides and fungicides.

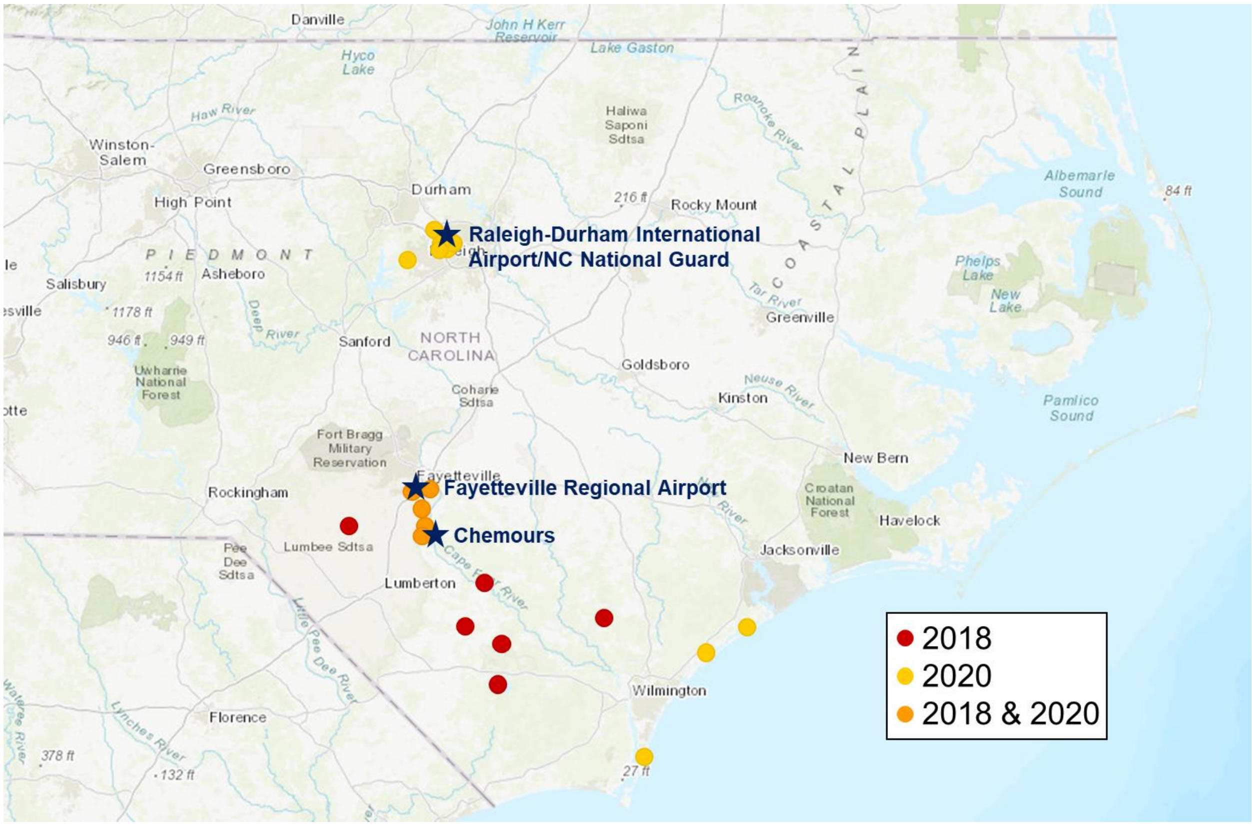


**Figure S1: Pine field sampling locations.** Colored by year(s) of sampling with points of interest noted.

**Table S1: Pine field sampling locations**

| Site Designation | Latitude, Longitude (degrees) | County | Year(s) Sampled | Needles Collected |
|---|---|---|---|---|
| **1** | 34.441, -78.530 | Bladen | 2018 | Tree/Shed |
| **2** | 34.533, -78.072 | Pender | 2018 | Tree |
| **3** | 34.506, -78.668 | Bladen | 2018 | Tree/Shed |
| **4** | 34.661, -78.612 | Bladen | 2018 | Tree/Shed |
| **5** | 34.836, -78.855 | Bladen | 2018, 2020 | Tree/Shed |
| **6** | 34.929, -78.856 | Cumberland | 2018, 2020 | Tree/Shed |
| **7** | 34.984, -78.893 | Cumberland | 2018, 2020 | Tree/Shed |
| **8** | 34.868, -78.853 | Cumberland | 2018, 2020 | Tree/Shed |
| **9** | 34.996, -78.832 | Cumberland | 2018, 2020 | Tree/Shed |
| **10** | 34.859, -79.189 | Robeson | 2018 | Tree |
| **11** | 34.294, -78.549 | Columbus | 2018 | Tree |
| **12** | 34.412, -77.640 | Pender | 2020 | Tree |
| **13** | 34.047, -77.921 | New Hanover | 2020 | Tree |
| **14** | 34.509, -77.463 | Pender | 2020 | Tree |
| **15** | 35.854, -78.795 | Wake | 2020 | Tree/Shed |
| **16** | 35.868, -78.782 | Wake | 2020 | Tree/Shed |

| | | | | |
|---|---|---|---|---|
| **17** | 35.859, -78.786 | Wake | 2020 | Tree/Shed |
| **18** | 35.877, -78.807 | Wake | 2020 | Tree/Shed |
| **19** | 35.836, -78.760 | Wake | 2020 | Tree/Shed |
| **20** | 35.787, -78.923 | Wake | 2020 | Tree/Shed |

**Table S2: Archived pine sample set**

| Specimen ID | Year | County |
|---|---|---|
| DUKE10008135 | 1938 | Durham |
| NCSC00034607 | 1947 | Carteret |
| DUKE10008133 | 1957 | Cumberland |
| DUKE10008159 | 1961 | Onslow |
| NCSC00034630 | 1967 | Wayne |
| DUKE10008158 | 1970 | Onslow |
| NCSC00034623 | 1977 | Onslow |
| DUKE10008136 | 1981 | Durham |
| DUKE10008118 | 1983 | Brunswick |
| DUKE10008160 | 1989 | Onslow |
| DUKE10008131 | 1991 | Cumberland |
| DUKE10008162 | 1995 | Robeson |
| DUKE10008110 | 1998 | Brunswick |
| DUKE10008130 | 2001 | Cumberland |
| DUKE10008138 | 2005 | Durham |

Prior to extraction, the randomized and blinded needles were homogenized using a methanol-washed stainless steel electric grinder. For field or archived samples, 2 g or 1 g (dry weight), respectively, of each sample homogenate was weighed into a 50-mL polypropylene tube. Five archived samples did not have enough material for 1 g, in which case 500 mg of sample homogenate was used. Aliquots of 15 mL methanol and 4 ng internal standards were mixed into the needle homogenates by vortexing for 15 seconds then sonicating for 30 minutes. Excess solid was allowed to separate for 15 minutes, then the supernatant was transferred to a fresh 50-mL polypropylene tube. The extraction steps were repeated with 10 mL of methanol, then the combined supernatants were vortexed for 5 seconds and filtered through a 0.45 µm syringe filter into a fresh 50-mL polypropylene tube. The extract was diluted with 25 mL of water, inverted 5 times and vortexed for 15 seconds.

Solid-phase extraction cleanup was performed using an Oasis WAX cartridge (Waters; Bedford, CA) which was conditioned with 4 mL of 0.1% ammonium hydroxide in methanol, 4 mL of methanol, and 4 mL of water. Extracts were passed through the cartridge at one drip per second. The cartridge was dried under vacuum for one minute then washed with 4 mL of acetate buffer (25 mM, pH 4) and 4 mL of methanol. Elution was performed using 4 mL of 0.1% ammonium hydroxide in methanol. The resulting eluents were collected in polypropylene tubes and dried under vacuum. Then, 200 µL of 2 mM ammonium acetate in 40:60 methanol/water was added. The reconstituted extracts were transferred to autosampler vials and stored at -20°C until analysis.

LC-IMS-MS Instrumental Analysis

Sample analyses were carried out on an Agilent 6560 IMS-qTOF instrument (Agilent Technologies; Santa Clara, CA) coupled with an Agilent 1290 Infinity LC system (Agilent Technologies; Santa Clara, CA) using a previously described method (*22*). Chromatographic separation of samples (2 µL injections) was performed using a C18 Agilent ZORBAX Eclipse Plus column (2.1 x 50 mm, 1.8 µm) with the gradient summarized in **Table S3** with a flow rate of 0.4 mL/min. Mobile Phase A was comprised of 5 mM ammonium acetate in water and Mobile Phase B was comprised of 5 mM ammonium acetate in 95:5 methanol/water. An Agilent Jet Stream ESI source (Agilent Technologies; Santa Clara, CA) was operated in negative ionization mode with the source conditions summarized in **Table S4**. IMS-MS settings are summarized in **Table S5**. Agilent ESI tune mix solution (Agilent Technologies; Santa Clara, CA) was directly injected to calibrate the instrument and calculate collision cross section (CCS) values for the PFAS analytes using a previously described and validated single-field calibration method (*22*). In short, tune mix ions with known CCS values served as calibrants for relating measured analyte drift times to CCS values.

**Table S3: LC gradient settings**

| Time (min) | % B |
|---|---|
| 0 | 10 |
| 0.5 | 10 |
| 2 | 30 |
| 14 | 95 |
| 14.5 | 100 |
| 16.5 (Stop Time) | 100 |
| 6 (Post Time) | 10 |

**Table S4: Ionization source settings**

| Parameter | Value | Units |
|---|---|---|
| Gas Temperature | 230 | ºC |
| Drying Gas | 11 | L/min |
| Nebulizer | 15 | psi |
| Sheath Gas Temperature | 350 | ºC |
| Sheath Gas Flow | 12 | L/min |
| $V_{cap}$ | 2500 | V |
| Nozzle Voltage | 0 | V |

**Table S5: IMS-MS settings**

| Parameter | Value | Units |
|---|---|---|
| Mass Range | 50-1700 | m/z |
| Trap Fill Time | 3900 | µs |
| Trap Release Time | 100 | µs |
| Frame Rate | 0.9 | Frames/s |
| IM Transient Rate | 16 | IM Transients/Frame |
| Max Drift Time | 60 | ms |
| TOF Transient Rate | 600 | Transients/IM Transients |
| Multiplexing Pulsing Sequence Length | 4 | bit |
| Drift Tube Entrance | -1574 | V |
| Drift Tube Exit | -224 | V |
| Rear Funnel Entrance | -217.5 | V |
| Rear Funnel Exit | -45 | V |

Quality Assurance and Data Analysis

Pooled quality control (QC) samples comprised of equal amounts of each pine sample homogenate and extraction blanks were prepared and analyzed alongside experimental samples. Additionally, Agilent ESI tune mix solution was utilized as an instrumental blank to monitor LC-IMS-MS instrument performance and confirm the absence of carryover approximately every 10-15 injections.

Each data file was demultiplexed using PNNL PreProcessor (v2020.03.23) with a signal intensity threshold of 20 counts. The data files were then single-field calibrated using Agilent ESI tune mix data collected in the same worklist as the sample using Agilent IM-MS Browser 10.0 software to relate measured drift times to associated CCS values.

Data was processed using Skyline-daily (v21.0.9.118) software with an in-house library containing PFAS class, name, molecular formula, adduct, *m/z*, retention time, and CCS values for >100 individual PFAS species ascertained from chemical standards. The PFAS compounds detected in at least one pine extract sample are listed in **Table S6**. Drift time filtering was used with a resolving power of 40 to eliminate noise due to the complex matrix. Targets that were not present in any samples were removed from the Skyline document.

**Table S6: Detected PFAS LC-IMS-MS parameters**

| Class | PFAS Analyte | Molecular Formula | *m/z* | Retention Time (min) | CCS (Å$^2$) |
|---|---|---|---|---|---|
| PFSA | PFDS | C10HF21O3S | 598.9238 | 11.8 | 186.2 |
| | PFNS | C9HF19SO3 | 548.9270 | 11.1 | 177.2 |
| | PFOS | C8HF17O3S | 498.9302 | 10.4 | 168.3 |
| | PFHpS | C7HF15O3S | 448.9334 | 9.2 | 159.2 |
| | PFHxS | C6HF13O3S | 398.9366 | 8.4 | 150.5 |
| | PFPeS | C5HF11O3S | 348.9398 | 7.0 | 142.2 |
| | PFBS | C4HF9O3S | 298.9430 | 5.2 | 133.6 |
| | PFPrS | C3HF7O3S | 248.9462 | 3.2 | 125.1 |
| | PFEtS | C2HF5O3S | 198.9494 | 1.3 | 117.4 |
| | TFMS | CF3SO3H | 148.9526 | 0.4 | 109.6 |
| | PFECHS | C8F15HO3S | 460.9334 | 9.0 | 153.3 |
| | 8Cl-PFOS | C8HClF16O3S | 514.9007 | 10.7 | 173.3 |
| | 1-MHpS | C8HF17O3S | 498.9302 | 10.4 | 163.2 |
| | m-PFOS | C8HF17O3S | 498.9302 | 10.1 | 164.9 |
| | dm-PFOS | C8HF17O3S | 498.9302 | 9.8 | 162.2 |
| | br-PFHpS | C7HF15O3S | 448.9334 | 9.3 | 123.1 |
| | br-PFHxS | C6HF13O3S | 398.9366 | 8.2 | 148.5 |
| | br-PFPeS | C5HF11O3S | 348.9398 | 6.8 | 140.2 |
| | br-PFBS | C4HF9O3S | 298.9430 | 5.0 | 131.6 |
| PFCA | PFHxDA | C16HF31O2 | 768.9510 | 14.0 | 203.7 |
| | PFTeDA | C14HF27O2 | 668.9574 | 13.3 | 187.6 |
| | PFTrDA | C13HF25O2 | 618.9606 | 12.9 | 179.7 |
| | PFDoA | C12HF23O2 | 568.9638 | 12.4 | 171.5 |
| | PFUdA | C11HF21O2 | 518.9670 | 11.8 | 163.4 |
| | PFDA | C10HF19O2 | 468.9702 | 11.1 | 155.3 |
| | PFNA | C9HF17O2 | 418.9734 | 10.3 | 147.0 |
| | PFOA | C8HF15O2 | 368.9766 | 9.4 | 139.5 |
| | PFHpA | C7HF13O2 | 318.9798 | 8.2 | 132.4 |
| | PFHxA | C6HF11O2 | 268.9830 | 6.7 | 125.1 |
| | PFPeA | C5HF9O2 | 218.9862 | 4.5 | 117.4 |
| | PFBA | C4HF7O2 | 168.9894 | 2.3 | 110.8 |
| | PFPrA | C3HF5O2 | 118.9926 | 10.5 | 119.3 |
| | br-PFOA | C8HF15O2 | 368.9766 | 9.3 | 136.5 |
| | br-PFHxA | C6HF11O2 | 268.9830 | 6.5 | 123.1 |
| | br-PFPeA | C5HF9O2 | 218.9862 | 4.2 | 116.4 |
| FT | 10:2 FTS | C12H5F21SO3 | 626.9551 | 12.4 | 203.6 |
| | 8:2 FTS | C10H5F17SO3 | 526.9615 | 11.1 | 185.8 |
| | 6:2 FTS | C8H5F13SO3 | 426.9679 | 9.3 | 168.1 |
| | 4:2 FTS | C6H5F9SO3 | 326.9743 | 6.5 | 150.4 |
| | 8:1 FTOH* | C9H3F17O | 448.9840 | 3.7 | 183.8 |
| PFE | R-EVE | C8H2F12O5 | 404.9632 | 2.4 | 146.3 |

| | | | | | |
|---|---|---|---|---|---|
| | HFPO-DA (GenX) | C6HF11O3 | 284.9779 | 7.1 | 127.5 |
| | PEPA | C5HF9O3 | 234.9811 | 5.0 | 120.9 |
| | PMPA | C4HF7O3 | 184.9843 | 2.6 | 115.2 |
| | PFMPA | C4HF7O3 | 184.9843 | 10.3 | 115.5 |
| | PFMOAA* | C3HF5O3 | 134.9875 | 5.2 | 108.0 |
| **PFESA** | Nafion byproduct 1 | C7HF13O5S | 442.9264 | 9.1 | 156.2 |
| | Nafion byproduct 2 | C7H2F14O5S | 462.9327 | 8.5 | 155.2 |
| | Nafion byproduct 4 | C7H2F12O6S | 440.9308 | 2.8 | 151.3 |
| | Nafion byproduct 6 | C6H2F12O4S | 396.9409 | 8.6 | 146.2 |
| | NVHOS | C4H2F8O4S | 296.9473 | 3.4 | 132.9 |
| | 9Cl-PF3ONS | C8HF16SO4Cl | 530.8956 | 10.8 | 170.2 |
| | 11Cl-PF3OUdS | C10HF20SO4Cl | 630.8892 | 12.1 | 170.2 |
| **PFASA** | NMeFOSE | C11H8F17NO3S | 555.9881 | 12.9 | 193.0 |
| | FOSAA | C10H4F17NO4S | 555.9517 | 10.5 | 181.7 |
| | FHxSA | C6H2F13SO2N | 397.9526 | 9.8 | 152.0 |
| | FBSAA | C6H4F9NO4S | 355.9645 | 6.9 | 147.2 |
| | FBSA | C4H2F9SO2N | 297.9590 | 6.7 | 135.0 |
| **PFP** | PFDPA | C10H2F21O3P | 598.9333 | 10.3 | 188.1 |
| | 6:2 diPAP | C16H9F26O4P | 788.9751 | 13.4 | 233.9 |
| | 6:2/8:2 diPAP | C18H9F30O4P | 888.9687 | 13.5 | 247.1 |
| **Unknown** | Unknown 248 | - | 247.9606 | 4.2 | 126.6 |
| | Unknown 265 | - | 264.9964 | 1.0 | 143.1 |
| | Unknown 281 | - | 280.9510 | 4.2 | 130.5 |
| | Unknown 297 | - | 296.9429 | 0.5 | 132.7 |
| | Unknown 336 | - | 335.9759 | 4.2 | 145.8 |
| | Unknown 347 | - | 346.9402 | 6.7 | 143.9 |
| | Unknown 348 | - | 347.9535 | 7.9 | 142.9 |
| | Unknown 436 | - | 435.9694 | 7.9 | 163.5 |
| | Unknown 486 | - | 485.9643 | 9.2 | 172.8 |
| | Unknown 489 | - | 488.9580 | 4.9 | 157.3 |
| | Unknown 571 | - | 570.9530 | 6.7 | 169.9 |
| | Unknown 589 | - | 588.9460 | 7.5 | 170.5 |
| | Unknown 659a | - | 658.9644 | 6.8 | 203.0 |
| | Unknown 659b | - | 658.9335 | 7.3 | 199.0 |

Annotations were based on mass accuracy, retention time alignment, CCS matching via drift time filtering, as well as co-elution and drift time alignment with labeled internal standards (if applicable). Additionally, tentatively identified and unknown targets were added to the Skyline document (**Table S7**). Tentative identifications were based on *m/z* (mass defect) and class-specific CCS versus *m/z* trends (*22*). Extracted ion intensities were exported to Excel for further analyses.

**Table S7: Internal standards used for signal normalization or estimating semi-quantitative levels**

| Native PFAS | Internal Standard | |
|---|---|---|
| | Field Samples | Temporal/Archived Samples |
| PFDS | $^{13}C_8$-PFOS | $^{13}C_8$-PFOS |
| PFNS | $^{13}C_8$-PFOS | $^{13}C_8$-PFOS |
| PFOS | $^{13}C_8$-PFOS | $^{13}C_8$-PFOS |
| PFHpS | $^{13}C_3$-PFHxS | $^{13}C_3$-PFHxS |
| PFHxS | $^{13}C_3$-PFHxS | $^{13}C_3$-PFHxS |
| PFPeS | $^{13}C_3$-PFHxS | $^{13}C_3$-PFHxS |
| PFBS | $^{13}C_3$-PFBS | $^{13}C_3$-PFBS |
| PFPrS | $^{13}C_3$-PFBS | $^{13}C_3$-PFBS |
| PFEtS | $^{13}C_3$-PFBS | $^{13}C_3$-PFBS |
| TFMS | $^{13}C_3$-PFBS | $^{13}C_3$-PFBS |
| PFecHS | $^{13}C_3$-PFHxS | $^{13}C_3$-PFHxS |
| 8Cl-PFOS | $^{13}C_8$-PFOS | $^{13}C_8$-PFOS |
| 1-MHpS | $^{13}C_8$-PFOS | $^{13}C_8$-PFOS |
| m-PFOS | $^{13}C_8$-PFOS | $^{13}C_8$-PFOS |
| dm-PFOS | $^{13}C_8$-PFOS | $^{13}C_8$-PFOS |
| br-PFHpS | $^{13}C_3$-PFHxS | $^{13}C_3$-PFHxS |
| br-PFHxS | $^{13}C_3$-PFHxS | $^{13}C_3$-PFHxS |
| br-PFPeS | $^{13}C_3$-PFHxS | $^{13}C_3$-PFHxS |
| br-PFBS | $^{13}C_3$-PFBS | $^{13}C_3$-PFBS |
| PFHxDA | $^{13}C_2$-PFTeDA | $^{13}C_2$-PFTeDA |
| PFTeDA | $^{13}C_2$-PFTeDA | $^{13}C_2$-PFTeDA |
| PFTrDA | $^{13}C_2$-PFTeDA | $^{13}C_2$-PFTeDA |
| PFDoA | $^{13}C_2$-PFDoA | $^{13}C_2$-PFDoA |
| PFUdA | $^{13}C_7$-PFUdA | $^{13}C_7$-PFUdA |
| PFDA | $^{13}C_6$-PFDA | $^{13}C_6$-PFDA |
| PFNA | $^{13}C_9$-PFNA | $^{13}C_9$-PFNA |
| PFOA | $^{13}C_8$-PFOA | $^{13}C_8$-PFOA |
| PFHpA | $^{13}C_4$-PFHpA | $^{13}C_4$-PFHpA |
| PFHxA | $^{13}C_5$-PFHxA | $^{13}C_5$-PFHxA |
| PFPeA | $^{13}C_5$-PFPeA | $^{13}C_5$-PFHxA |
| PFBA | $^{13}C_5$-PFHxA | $^{13}C_5$-PFHxA |
| PFPrA | $^{13}C_9$-PFNA | $^{13}C_9$-PFNA |
| br-PFOA | $^{13}C_8$-PFOA | $^{13}C_8$-PFOA |
| br-PFHxA | $^{13}C_5$-PFHxA | $^{13}C_5$-PFHxA |
| br-PFPeA | $^{13}C_5$-PFPeA | $^{13}C_5$-PFPeA |
| 10:2 FTS | $^{13}C_8$-PFOS | $^{13}C_8$-PFOS |
| 8:2 FTS | $^{13}C_8$-PFOS | $^{13}C_8$-PFOS |
| 6:2 FTS | $^{13}C_3$-PFHxS | $^{13}C_3$-PFHxS |

| | | |
|---|---|---|
| 4:2 FTS | $^{13}C_3$-PFBS | $^{13}C_3$-PFBS |
| 8:1 FTOH | $^{13}C_3$-PFBS | $^{13}C_3$-PFBS |
| R-EVE | $^{13}C_5$-PFHxA | $^{13}C_3$-GenX |
| HFPO-DA (GenX) | $^{13}C_5$-PFHxA | $^{13}C_3$-GenX |
| PEPA | $^{13}C_5$-PFHxA | $^{13}C_3$-GenX |
| PMPA | $^{13}C_5$-PFHxA | $^{13}C_3$-GenX |
| PFMPA | $^{13}C_9$-PFNA | $^{13}C_3$-GenX |
| PFMOAA | $^{13}C_5$-PFHxA | $^{13}C_3$-GenX |
| Nafion byproduct 1 | $^{13}C_3$-PFHxS | $^{13}C_3$-PFHxS |
| Nafion byproduct 2 | $^{13}C_3$-PFHxS | $^{13}C_3$-PFHxS |
| Nafion byproduct 4 | $^{13}C_3$-PFBS | $^{13}C_3$-PFBS |
| Nafion byproduct 6 | $^{13}C_3$-PFHxS | $^{13}C_3$-PFHxS |
| NVHOS | $^{13}C_3$-PFBS | $^{13}C_3$-PFBS |
| 9Cl-PF3ONS | $^{13}C_8$-PFOS | $^{13}C_8$-PFOS |
| 11Cl-PF3OUdS | $^{13}C_8$-PFOS | $^{13}C_8$-PFOS |
| NMeFOSE | $^{13}C_8$-PFOS | $^{13}C_8$-PFOS |
| FOSAA | $^{13}C_8$-PFOS | $^{13}C_8$-PFOS |
| FHxSA | $^{13}C_3$-PFHxS | $^{13}C_3$-PFHxS |
| FBSAA | $^{13}C_3$-PFHxS | $^{13}C_3$-PFHxS |
| FBSA | $^{13}C_3$-PFBS | $^{13}C_3$-PFBS |
| PFDPA | $^{13}C_9$-PFNA | $^{13}C_9$-PFNA |
| 6:2 diPAP | $^{13}C_2$-PFTeDA | $^{13}C_2$-PFTeDA |
| 6:2/8:2 diPAP | $^{13}C_2$-PFTeDA | $^{13}C_2$-PFTeDA |
| Unknown 248 | $^{13}C_3$-PFBS | $^{13}C_3$-PFBS |
| Unknown 265 | $^{13}C_3$-PFBS | $^{13}C_3$-PFBS |
| Unknown 281 | $^{13}C_3$-PFBS | $^{13}C_3$-PFBS |
| Unknown 297 | $^{13}C_3$-PFBS | $^{13}C_3$-PFBS |
| Unknown 336 | $^{13}C_3$-PFBS | $^{13}C_3$-PFBS |
| Unknown 347 | $^{13}C_5$-PFHxA | $^{13}C_5$-PFHxA |
| Unknown 348 | $^{13}C_4$-PFHpA | $^{13}C_4$-PFHpA |
| Unknown 436 | $^{13}C_4$-PFHpA | $^{13}C_4$-PFHpA |
| Unknown 486 | $^{13}C_8$-PFOA | $^{13}C_8$-PFOA |
| Unknown 489 | $^{13}C_3$-PFBS | $^{13}C_3$-PFBS |
| Unknown 571 | $^{13}C_5$-PFHxA | $^{13}C_5$-PFHxA |
| Unknown 589 | $^{13}C_5$-PFHxA | $^{13}C_5$-PFHxA |
| Unknown 659a | $^{13}C_5$-PFHxA | $^{13}C_5$-PFHxA |
| Unknown 659b | $^{13}C_5$-PFHxA | $^{13}C_5$-PFHxA |

Peak areas of analytes with labeled internal standards were normalized to the respective internal standard, and those without matching internal standards were normalized to surrogate standards, which were based on structural similarity and retention time proximity (**Table S6**). Normalized peak areas of analytes detected in extraction blanks were subtracted from the sample normalized peak areas to assess only the signal extracted from the needles themselves. For the archived samples, three blanks were averaged and only signal outside of three standard deviations from the blank mean were considered. Semi-quantitative concentration estimates were made for analytes with labeled internal standards based on the light/heavy ratio, the amount of internal standard, and the amount of material used for extraction. Analytes without matching internal standards were related to the amount of material used for extraction (normalized peak area/g) but not quantified. These relative and semi-quantitative values were imported into the ToxPi GUI for visualization and further analyses. The interactive map, Spatial and Temporal ToxPi Profiles of Per- and Polyfluoroalkyl Substances (PFAS) in North Carolina Pine Needles, was created using the ToxPi*GIS Toolkit, which generates ToxPi feature layers that can be used from with ArcGIS. A walkthrough of the steps used to make this map are in the toolkit documentation, which is linked at www.toxpi.org.

### Supplementary Text

Qualitative and Quantitative Data

Extracted ion chromatograms for each PFAS target across all experimental samples, blanks, and pooled quality control samples are available via Skyline's online repository Panorama (https://panoramaweb.org/pfas_pine.url). Processed data for the 61 identified PFAS is also available as an interactive ToxPi*ArcGIS map (https://arcg.is/19vXuK0).

**Table S8: Archived pine concentration estimations**

| PFAS | 1967 W | 1970 O | 1977 O | 1981 D | 1983 B | 1989 O | 1991 C | 1995 R | 1998 B | 2001 C | 2005 D |
|---|---|---|---|---|---|---|---|---|---|---|---|
| **PFOS** | 33.55676 | 14.33264 | 227.7124 | 19.64127 | 17.83696 | 197.1817 | 8.88239 | 29.01167 | 34.92666 | 19.24914 | 5.746803 |
| **PFHxS** | 27.15817 | 13.21294 | 22.00576 | 4.451343 | 11.11259 | 35.89003 | ND | 4.92617 | 1.486732 | 16.3247 | 5.400338 |
| **PFBS** | ND | ND | ND | ND | ND | ND | ND | ND | ND | 0.543754 | ND |
| **PFTeDA** | 3773.371 | 468.4296 | 403.479 | 1326.89 | 2938.63 | ND | 2011.766 | 3010.678 | 1117.931 | 724.201 | 1896.128 |
| **PFDoA** | 141.8947 | ND | ND | 201.3709 | ND | ND | ND | 119.1291 | ND | 303.0915 | 11.17723 |
| **PFUdA** | ND | ND | ND | ND | ND | ND | ND | ND | ND | 0.111429 | 0.419157 |
| **PFDA** | 0.320177 | ND | ND | ND | 0.354995 | 10.43423 | ND | 1.491039 | ND | ND | 1.182514 |
| **PFNA** | ND | ND | ND | ND | 7.143599 | 10.57547 | ND | 5.363673 | 11.98998 | 5.893119 | ND |
| **PFOA** | ND | 54.13048 | ND | 7.416052 | ND | 332.7059 | 61.93348 | ND | 97.14829 | ND | ND |
| **PFHpA** | ND | ND | ND | ND | ND | ND | ND | ND | 8.654581 | ND | ND |

W = Wayne County, O = Onslow County, D = Durham County, B = Brunswick County, C = Cumberland County, R = Robeson County; ND = Not Detected. Values reported as pg/g dry weight.

### Table S9: Field sample concentration estimations

| Site ID | Year | PFTeDA | PFDoA | PFUdA | PFDA | PFNA | PFOA | PFHxA | PFHpA | PFPeA | PFBA | PFOS | PFHxS | PFBS |
|---|---|---|---|---|---|---|---|---|---|---|---|---|---|---|
| **1** | **2017** | 101.50 | 7.55 | 4.64 | 21.54 | 32.45 | 291.36 | ND | 283.69 | ND | ND | ND | ND | 10.77 |
| **1** | **2018** | 39.06 | 4.87 | 2.91 | 15.81 | 54.47 | 2001.95 | ND | 233.43 | ND | ND | ND | ND | 8.87 |
| **2** | **2018** | 7.12 | 8.59 | 4.72 | 27.72 | 67.16 | 950.28 | 22.84 | 353.15 | ND | ND | 100.48 | ND | 4.62 |
| **3** | **2017** | 53.60 | 9.08 | 0.86 | 21.43 | 40.94 | 428.64 | ND | 291.54 | ND | ND | 5.56 | ND | 0.76 |
| **3** | **2018** | 54.96 | 11.94 | 6.63 | 49.81 | 118.52 | 1363.52 | 173.65 | 603.26 | ND | ND | ND | ND | 8.88 |
| **4** | **2018** | 94.39 | 8.99 | ND | 2.83 | ND | 169.60 | ND | 149.38 | ND | ND | ND | ND | 15.65 |
| **4** | **2017** | 78.25 | 8.74 | 8.84 | 38.28 | 114.99 | 577.75 | ND | 631.23 | ND | ND | ND | ND | 18.23 |
| **5** | **2019** | ND | 5.07 | 109.17 | 16.16 | 39.05 | 207.44 | ND | 136.59 | ND | 5.16 | 3.34 | 18.33 | 40.94 |
| **5** | **2020** | ND | ND | 121.32 | 1.82 | 39.11 | 217.39 | 23.20 | 166.08 | ND | 20.41 | 1.61 | 21.87 | 22.17 |
| **5** | **2018** | 75.22 | 6.30 | 26.13 | 27.04 | 59.51 | 514.96 | 89.67 | 400.62 | ND | ND | 31.89 | ND | 42.32 |
| **5** | **2017** | 42.80 | 9.62 | 5.10 | 26.70 | 88.39 | 645.82 | 185.84 | 496.12 | 86.97 | ND | 37.86 | ND | 84.40 |
| **6** | **2020** | ND | 19.08 | 2207.42 | 5.98 | ND | 84.26 | ND | 110.31 | ND | ND | ND | 39.65 | 14.70 |
| **6** | **2019** | ND | 0.83 | 138.57 | 4.02 | 28.80 | 121.65 | ND | 89.75 | ND | ND | 4.12 | 11.93 | 24.93 |
| **6** | **2017** | 40.22 | 5.53 | ND | 14.78 | 21.12 | 199.74 | 53.51 | 301.58 | ND | ND | ND | ND | 14.20 |
| **6** | **2018** | 20.72 | 1.80 | 8.65 | 18.37 | 34.05 | 381.40 | 70.34 | 330.64 | ND | ND | ND | ND | 17.68 |
| **7** | **2020** | ND | 21.60 | 914.80 | 12.42 | 44.47 | 182.15 | 67.14 | 131.80 | ND | ND | 11.17 | 135.48 | 33.97 |
| **7** | **2019** | ND | 1.07 | 135.71 | 4.77 | 10.76 | 36.77 | ND | ND | ND | ND | 66.95 | 54.23 | 37.98 |
| **7** | **2017** | 62.11 | 10.34 | 6.02 | ND | 4.88 | 67.50 | ND | 121.21 | ND | ND | ND | ND | 23.92 |
| **7** | **2018** | 34.97 | 4.05 | 0.72 | 11.75 | 13.38 | 246.71 | 117.83 | 289.69 | ND | ND | ND | ND | 15.54 |
| **8** | **2019** | ND | ND | 96.78 | 8.57 | 27.40 | 177.73 | 68.18 | 155.24 | ND | ND | 8.75 | 14.80 | 51.29 |
| **8** | **2020** | 123.22 | ND | 55.83 | 4.83 | 3.05 | 3.45 | 16.46 | ND | ND | ND | 6.75 | 30.58 | 31.82 |
| **8** | **2017** | 47.39 | ND | 5.13 | 5.43 | 26.97 | 253.11 | ND | 301.81 | ND | 12.69 | ND | ND | 45.41 |
| **8** | **2018** | 77.07 | 3.37 | 8.30 | 21.82 | 62.59 | 412.94 | 219.61 | 450.57 | ND | 14.28 | ND | ND | 56.62 |
| **9** | **2020** | ND | 3.98 | 681.91 | 4.82 | 28.07 | 154.43 | ND | 132.00 | ND | ND | 2.20 | 15.55 | 29.56 |
| **9** | **2019** | ND | 1.79 | 545.36 | 9.21 | 22.94 | 193.85 | 13.99 | 208.27 | ND | ND | 2.42 | 29.03 | 43.81 |
| **9** | **2018** | 73.44 | 10.39 | 7.35 | 24.26 | 6.50 | 98.48 | ND | 147.63 | ND | ND | ND | ND | 18.20 |
| **9** | **2017** | 62.48 | 16.72 | ND | 4.85 | 8.52 | 96.26 | ND | 98.23 | ND | ND | ND | ND | 37.78 |
| **10** | **2018** | 37.62 | 7.44 | 4.72 | 18.89 | 28.75 | 15038.89 | 101.21 | 286.51 | ND | ND | ND | ND | 4.42 |
| **11** | **2018** | 67.25 | 3.02 | ND | 8.54 | 7.14 | 13924.95 | ND | 221.87 | ND | ND | ND | ND | ND |
| **12** | **2020** | 37.06 | 0.50 | 3.26 | 14.14 | 26.07 | 190.36 | 32.74 | 210.64 | ND | ND | 4.44 | ND | ND |
| **13** | **2020** | 27.52 | 0.38 | 5.84 | 20.02 | 3.31 | 72.93 | 13.30 | 136.17 | ND | ND | 36.80 | ND | 49.99 |
| **14** | **2020** | 80.03 | 2.34 | 6.52 | 33.67 | 67.30 | 370.61 | 2.79 | 331.71 | ND | ND | 9.08 | ND | 17.34 |
| **15** | **2019** | 143.71 | ND | 84.22 | 6.38 | 7.09 | 55.33 | ND | 57.42 | ND | ND | 5.16 | 23.22 | 12.09 |
| **15** | **2020** | 57.97 | 15.72 | 187.78 | 23.49 | 29.67 | ND | ND | ND | ND | ND | 14.37 | 57.78 | 14.47 |
| **16** | **2019** | ND | 3.71 | 485.91 | 7.54 | 49.02 | 187.96 | 2635.63 | 368.97 | 31623.52 | 26.27 | 204.05 | 8306.69 | 6672.06 |
| **16** | **2020** | ND | 7.19 | 875.32 | 18.80 | 64.17 | 579.43 | 23970.72 | 2669.00 | 189646.89 | 808.93 | 2076.03 | 26822.01 | 23379.76 |
| **17** | **2019** | ND | ND | 554.43 | 11.75 | 35.03 | 210.92 | ND | 173.04 | ND | ND | 8.89 | 54.87 | 35.98 |
| **17** | **2020** | ND | ND | 170.54 | 14.15 | 35.99 | 158.81 | 17.19 | 111.41 | ND | ND | 4.31 | 15.04 | 5.97 |
| **18** | **2019** | ND | ND | 196.47 | 15.37 | 36.28 | 172.24 | ND | 103.74 | ND | ND | 14.93 | 29.40 | 39.44 |
| **18** | **2020** | ND | 5.11 | 97.45 | 12.64 | 35.30 | 189.67 | 55.78 | 151.39 | ND | ND | 5.07 | 12.56 | 8.56 |
| **19** | **2020** | ND | 19.87 | 930.73 | 19.43 | 69.71 | 194.54 | 44.91 | 168.95 | ND | ND | 4.40 | 22.75 | 6.13 |

| | | | | | | | | | | | | | | |
|---|---|---|---|---|---|---|---|---|---|---|---|---|---|---|
| **19** | **2019** | ND | 2.51 | 65.51 | 3.46 | 15.64 | 4.81 | ND | 56.23 | ND | ND | 11.27 | 23.81 | 11.41 |
| **20** | **2020** | ND | 5.18 | 1217.30 | 24.30 | 51.98 | 209.60 | ND | 179.18 | ND | ND | 3.13 | 42.26 | 12.28 |
| **20** | **2019** | ND | 2.66 | 83.13 | 21.23 | 31.97 | 129.02 | ND | 72.75 | ND | ND | 14.57 | 20.53 | 40.14 |

ND = Not Detected. Field sample location IDs correspond to those listed in **Table S1**. Values reported as pg/g dry weight.

Multidimensional Data and Isomers

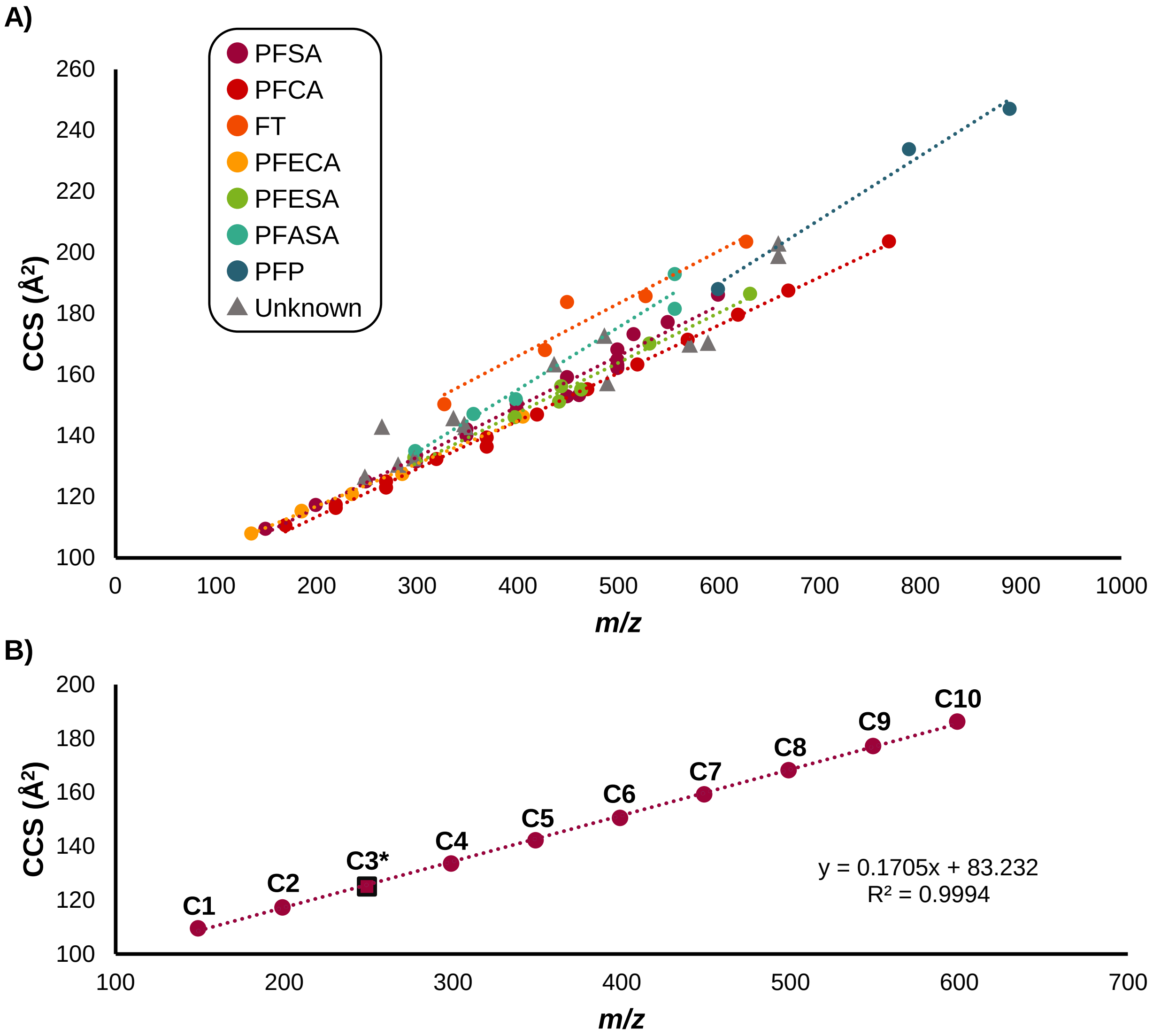


**Figure S2: IMS-MS CCS versus *m/z* trendlines.** (**A**) CCS versus *m/z* trendlines for the PFAS analyzed colored by subclass. The PFAS headgroups contribute to the observed CCS for a given *m/z* in each subclass. Some subclasses have multiple headgroups or differing chemistries such as chlorinated replacements and cyclic compounds which causes the imperfect fit in some cases. Unknown PFAS are also displayed to demonstrate that they fall within the expected CCS and *m/z* window. (**B**) CCS versus *m/z* trendline for the PFSA homologous series, i.e. varying chain lengths of $CF_2$ with the same sulfonic acid headgroup. The carbon numbers are labeled as CX, where X is the number of carbons. For example, C8 is perfluorooctanesulfonic acid (PFOS). The 3-carbon PFSA (C3*) was not present in the standard library used to annotate the data, but was able to be confidently identified due to the homologous series trendline.

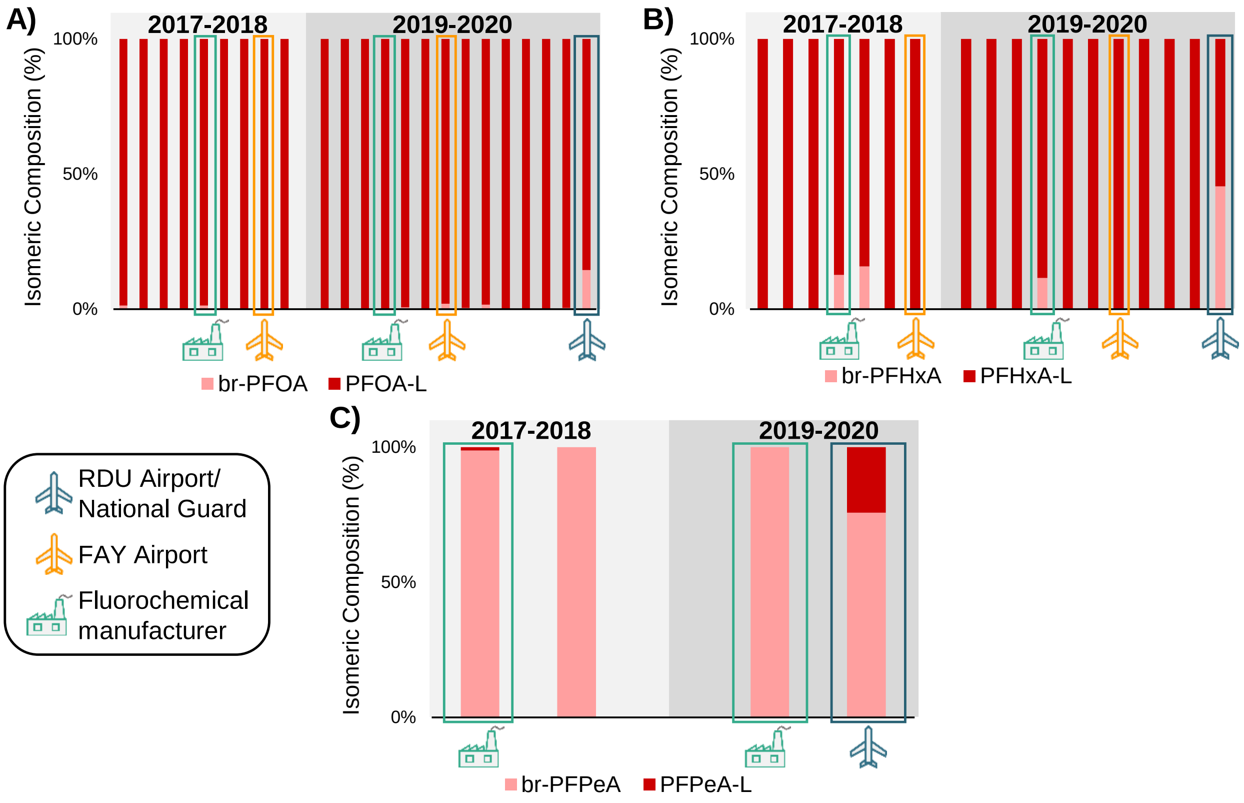


**Figure S3: PFCA isomeric composition**. Percent composition of all detected branched and linear PFCA isomer pairs. Sampling sites are ordered by time (2017-2018 and 2019-2020) and location left to right from southeastern NC to central NC. Discussed PFAS point sources including the RDU International Airport/NC National Guard facility, FAY Regional Airport, and a fluorochemical manufacturer are marked. Only samples where at least one of the two isomers were detected are included. PFCA isomer pairs include branched and linear (**A**) PFOA (8-carbon), (**B**) PFHxA (6-carbon), and (**C**) PFPeA (5-carbon).

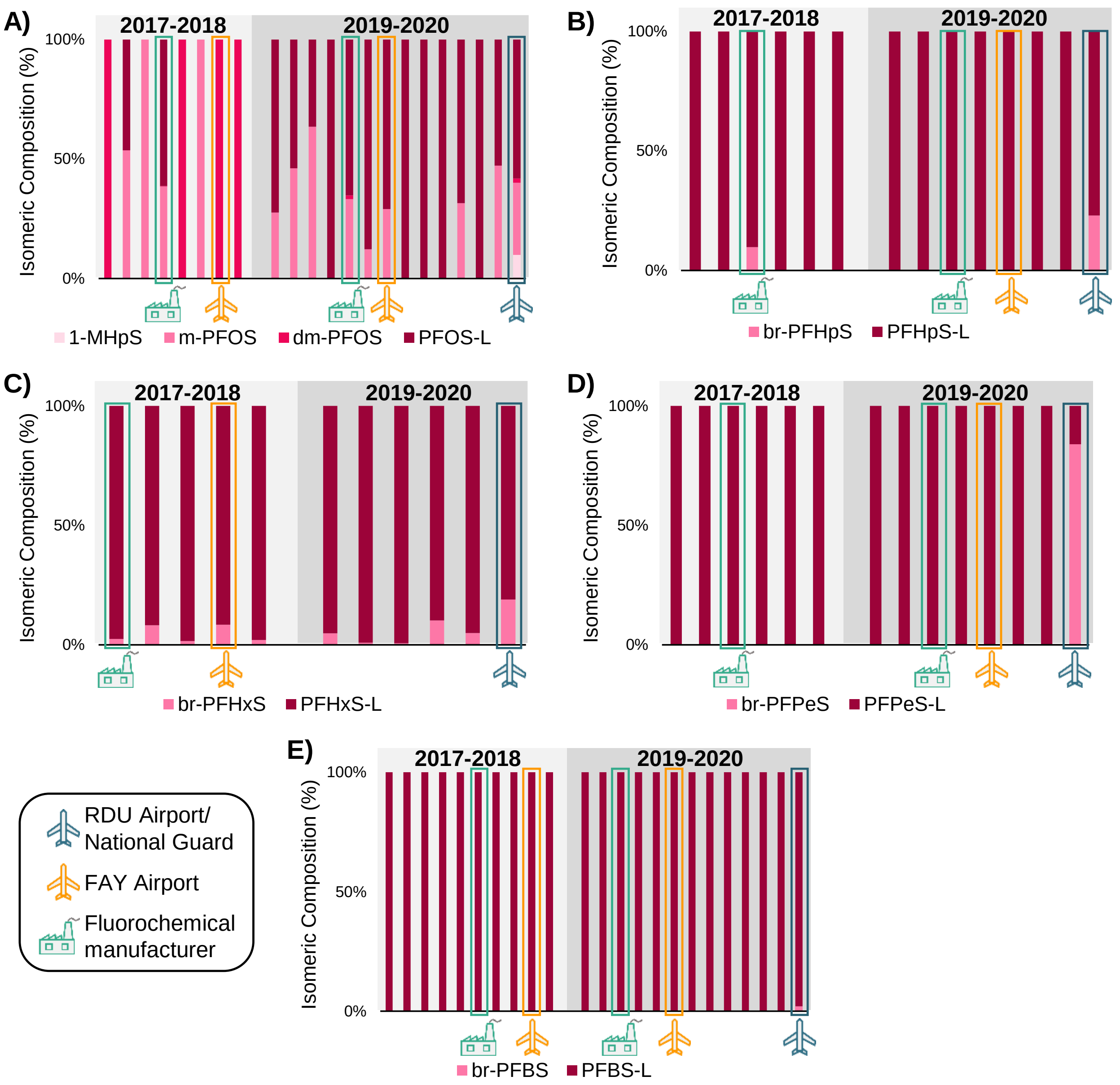


**Figure S4: PFSA isomeric composition**. Percent composition of all detected branched and linear PFSA isomers. Sampling sites are ordered by time (2017-2018 and 2019-2020) and location left to right from southeastern NC to central NC. Discussed PFAS point sources including the RDU International Airport/NC National Guard facility, FAY Regional Airport, and a fluorochemical manufacturer are marked. Only samples where at least one of the isomers were detected are included. PFSA isomers include (**A**) linear, dimethylated, and methylated PFOS (8-carbon) as well as the methylated 1-MHpS isomer, and linear and branched (**B**) PFHpS (7-carbon), (**C**) PFHxS (6-carbon), (**D**) PFPeS (5-carbon), and (**E**) PFBS (4-carbon).

Point Sources

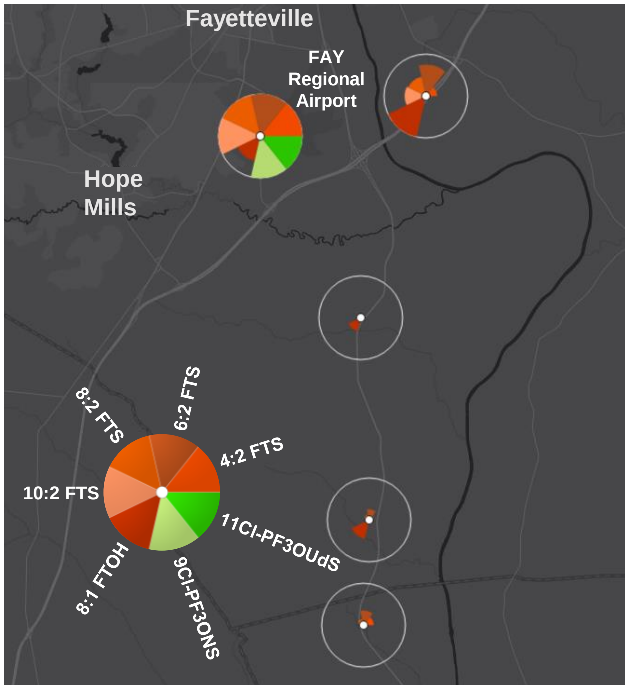


**Figure S5: FAY Regional Airport trends.** Map of southern Fayetteville, NC with averaged fluorotelomer and chlorofluoroethers ToxPi profiles of pine needles collected from 5 locations in 2017-2020. The peak levels of each of these compounds were detected at one of the two sites nearest the FAY Regional Airport. The location nearest the FAY Regional Airport (0.5 mi) had the highest levels of each fluorotelomer sulfonate, which ranged from 2-8X higher than the study average over the same time period. Additionally, this location was the only one at which the two chlorofluoroethers, 9Cl-PF3ONS and 11-ClPF3OUdS, were detected.
